# Antibody repertoire analysis of tumor-infiltrating B cells reveals distinct signatures and distributions across tissues

**DOI:** 10.1101/2021.03.17.435773

**Authors:** Ligal Aizik, Yael Dror, David Taussig, Adi Barzel, Yaron Carmi, Yariv Wine

**Affiliations:** The Shmunis School of Biomedicine and Cancer Research, The George S. Wise Faculty of Life Sciences, Tel Aviv University, Tel Aviv, 69978, Israel; The School of Neurobiology, Biochemistry and Biophysics, The George S. Wise Faculty of Life Sciences, Tel Aviv University, Tel Aviv, 69978, Israel; Department of Pathology, Sackler School of Medicine, Tel Aviv University, Tel Aviv 69978, Israel

**Keywords:** Antibody repertoire, BCR-Seq, Tumor infiltrating Lymphocytes, TIL, Next generation Sequencing, VDJ recombination, B cell, Triple negative breast Cancer, TNBC, somatic hypermutation, class switch recombination

## Abstract

The role of B cells in the tumor microenvironment (TME) has largely been under investigated, and data regarding the antibody repertoire encoded by B cells in the TME and the adjacent lymphoid organs are scarce. Here, we utilized B cell receptor high-throughput sequencing (BCR-Seq) to profile the antibody repertoire signature of tumor-infiltrating lymphocyte B cells (TIL-Bs) in comparison to B cells from three anatomic compartments in a mouse model of triple-negative breast cancer. We found that TIL-Bs exhibit distinct antibody repertoire measures, including high clonal polarization and elevated somatic hypermutation rates, suggesting a local antigen-driven B-cell response. Importantly, TIL-Bs were highly mutated but non-class switched, suggesting that class-switch recombination may be inhibited in the TME. Tracing the distribution of TIL-B clones across various compartments indicated that they migrate to and from the TME. The data thus suggests that antibody repertoire signatures can serve as indicators for identifying tumor-reactive B cells.

## Introduction

B cells and T cells are distinct effector cells in the adaptive arm of the immune system; yet, they often act in synergy to eradicate pathological processes. For more than two decades, research on the immune response to tumors has focused mainly on T cells, with B cells being under investigated and their roles in the tumor microenvironment (TME) remaining controversial^1,2^. Recent studies have, however, revealed that B cells may play a crucial role in tumor immunity, as these cells consistently comprise a substantial cellular component of the TME. Indeed, it has been reported that tumor-infiltrating lymphocyte B cells (TIL-Bs) can reach up to 40% of all TILs in different types of cancer^3,4^. Moreover, several reports have described tertiary lymphoid structures (TLN) that contain a relatively high portion of TIL-Bs^5–7^.

Several possible functions have been attributed to TIL-Bs. Among these functions, they can serve as antibody-secreting cells, namely, as plasma cells producing antibodies that inhibit tumor growth by targeting tumor-associated antigens (TAAs) and inducing antibody-dependent cell cytotoxicity or complement-dependent cytotoxicity^8^. TIL-Bs can also function as antigen-presenting cells that trigger an anti-tumor T cell response^9,10^. Of particular importance, TIL-Bs can exhibit direct cytotoxic activity, killing tumor cells via the Fas/FasL pathway^11^: Activated B cells in tumor-draining lymph nodes (DLNs) have been shown to express the Fas ligand, to be upregulated upon engagement with cells of the 4T1 triplenegative breast cancer (TNBC) cell line, and, in turn, to exhibit cytotoxic activity^12^. The discovery of regulatory B cells that secrete cytokines suggests an additional possible role of B cells in the TME^13,14^ It has been reported, for example, that the frequency of interleukin (IL)-10-producing B cells in the TME of some cancer types is negatively correlated with that of interferon (IFN)-γ-producing CD8^+^ T cells but positively correlated with that of regulatory Foxp3^+^ CD4^+^ T cells. These data thus demonstrate a possible regulatory role for B cells in the TME^15^. The above research notwithstanding, reports describing TIL-B activity in cancer remain contradictory, with some studies claiming TIL-B activity to be immunosuppressive^16–18^ and others assigning protective roles to these cells^19–21^. These contradictory pieces of evidence have led cancer scientists to hold differing views about whether immunotherapies should be aimed to enhance or inhibit the activity of the B cells in the TME.

The analysis of antibodies that become anchored to the membrane of the B cell as B cell receptors (BCRs) was made possible by virtue of advances in the next-generation sequencing (NGS) of BCRs (BCR-Seq), which has revolutionized our understanding of B-cell-mediated immunity. This technology, coupled with the development of advance computational tools, has enabled the elucidation of the enormous antibody diversity, which, in turn, provides important information about the nature of the B cell response following a challenge^22^.

BCR-Seq enables the exploration of antibody repertoire measures that are associated with the unique diversification mechanisms that B cells undergo during their development. Initially, B cells in the bone marrow are subjected to a V(D)J chromosomal rearrangement process, followed by N/P non-templated nucleotide deletion/insertion^23^. Subsequently, in response to an antigenic challenge, B cells are clonally expanded and diversify further in a process known as affinity maturation, which includes somatic hypermutation (SHM), followed by clonal selection for B cells that encode productive high-affinity antibodies^24^. In addition, class-switch recombination (CSR), in which the antibody isotype is altered (e.g., IgM to IgG), contributes to the effector function that is attributed to antibodies. The abovementioned mechanisms facilitate the generation of an enormous diversity of antibodies that can exceed the theoretical diversity of 10^13^25^. Thus, repertoire measures, such as B cell clonal architecture, SHM rate and isotype distribution, can be exploited to facilitate our understanding of the antigen-driven B cell response in cancer^26–29^.

To provide new insights into the functional role of B cells in the TME, we performed BCR-Seq on B lymphocytes from four tissue types, namely, tumors, DLNs, blood, and bone marrow, derived from TNBC-induced mice. We tracked the antibody repertoire measures of the TIL-B clones compared to the measures of the B cell clones in the other three tissue types with the aim to identify the signatures that characterize B cells engaged with tumor cells and/or TAAs and to determine whether they are subjected to positive selection processes following an antigen-driven response in the TME and the adjacent DLNs. We found that the TIL-B compartment is dominated by a restricted number of highly expanded clones exhibiting high rates of SHM. Interestingly, these clones were mostly IgM^+^ clones, suggesting that the CSR of TIL-Bs is impaired. Moreover, a particular subset of TIL-Bs was found in all compartments, suggesting that this specific subset of B cells migrates from and to the TME.

## Results

### Experimental and computational study design

Tracking B cell clones, either in a temporal manner or between tissues, can be achieved by identifying their antibody variable regions and particularly their complementarity-determining region 3 (CDR3) of the heavy chain (CDRH3), which is used as a B cell clonal “barcode.” Since the CDRH3 is a product of the V(D)J rearrangement, it exhibits the highest diversity in the antibody heavy chain variable region (V_H_) and is thought to play a key role in antigen recognition^25,30,31^. Since it is generally accepted that the CSR from IgM to IgG represents progression of the immune response^32,33^, we thus focused our analysis on the antibody V_H_ of both the IgG and IgM isotypes. To study the repertoire measures of TIL-Bs and to track their clonal distribution across tissues, we established an experimental and computational platform for the analysis of the V_H_ sequences obtained from four mice treated with 4T1 cells (vs two untreated mice as control) (*Figure 1A*). At 23 days post tumor-cell inoculation, B cells were collected from four tissue types (tumor, blood, DLNs and bone marrow) and lysed for purification of total mRNA. Recovered mRNA was reverse transcribed to generate cDNA, and the V_H_ sequences were amplified by using a primer set specific for the V_H_ and the constant region 1 of the heavy chain (C_H_1) of the *IGHM* and *IGHG* genes (Supplementary *Table S1*). Generated V_H_ amplicons were prepared for multiplex sequencing on a MiSeq Illumina platform (2×300 bp) by the addition of adaptors and sample-specific barcodes. For the untreated mice, B cells from three tissue types were isolated and subjected to the same process as that for the treated mice.

**Figure 1.**
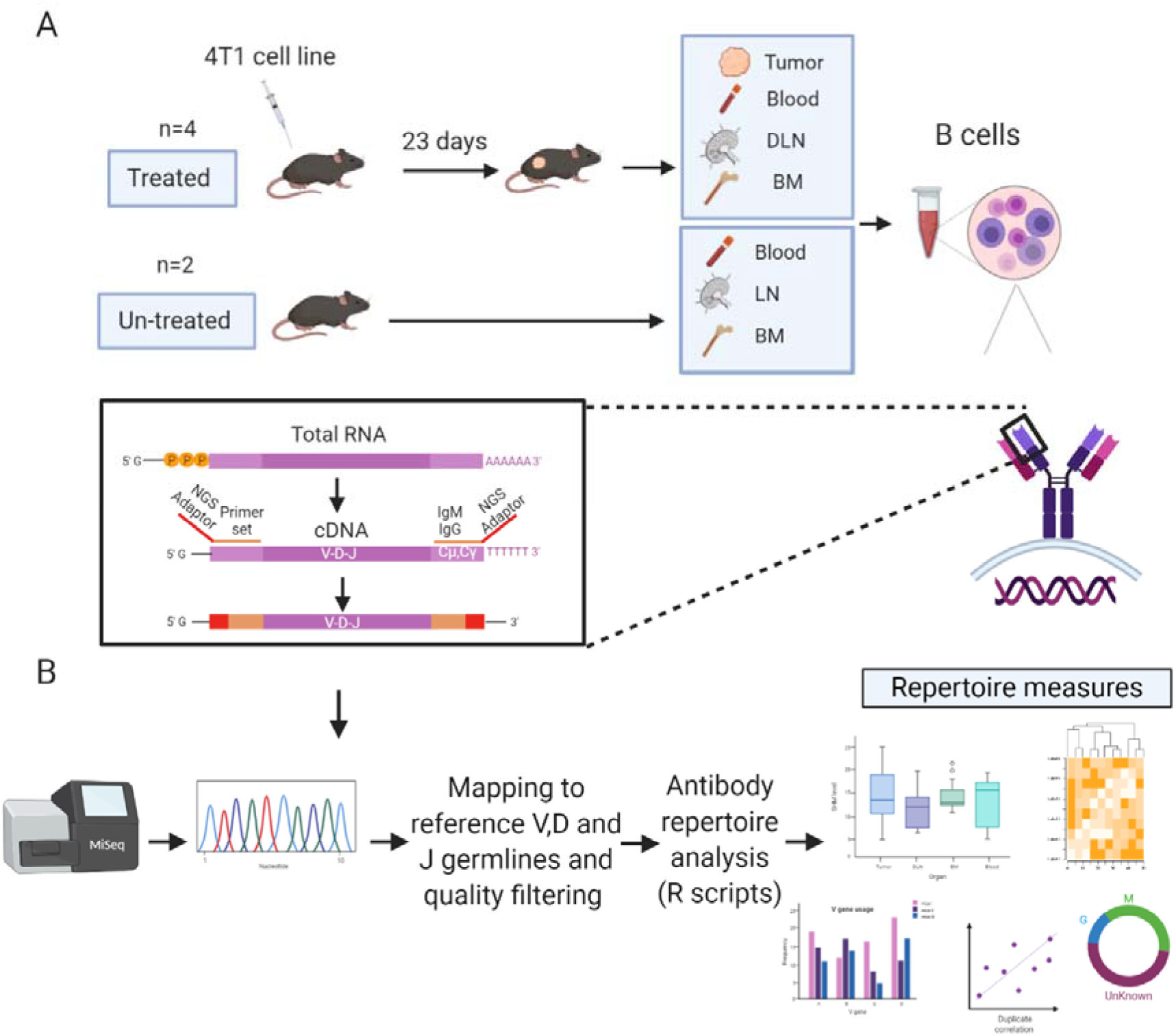
Experimental and computational platform for antibody repertoire analysis of TIL-Bs. (**A**) BALB/c female mice (6–8 weeks old, n = 4) were inoculated with 4T1 tumor cells into the mammary fat pad. On day 23, mice were sacrificed, and B cells (IgM/IgG/CD138+) were isolated from four tissue types: tumors, bone marrow, draining lymph nodes (DLNs), and peripheral blood. Recovered mRNA was used as a template to generate NGS libraries. The same experimental pipeline was applied to naïve mice (n = 2) as the control. (**B**) NGS libraries were sequenced using the Illumina MiSeq 2×300 platform, and the resultant paired-end antibody V_H_ sequences were assembled and aligned using MiXCR^34^. V_H_ sequences were then filtered and high-quality V_H_ sequence datasets were used to generate antibody repertoire measures (developed under R) of TIL-Bs and B cells from other tissues.

BCR-Seq of B cells from tissues of all four mice yielded a total of 5.3×10^7^ sequences. V_H_ sequences were paired-end aligned and annotated by alignment to reference germlines, as found in the ImMunoGeneTics server (IMGT^34^) using MiXCR^35^, to obtain annotation by regions including the frameworks (FRs), the CDRs, and the full-length VDJ region. Aligned V_H_ sequences were filtered according to chosen criteria (see Methods) before establishing the antibody sequence dataset to be used for generating the repertoire measures (*Figure 1B*).

### BCR-Seq analysis

To reduce the errors that are introduced during PCR amplification and sequencing, we carried out BCR-Seq on technical duplicates from each sample. Only V_H_ sequences that appeared in both replicates were considered valid. To estimate whether the sequencing depth was adequate, the BCR-Seq data for each sample was subjected to rarefaction analysis (a method used routinely in ecology to quantify species diversity^36^; see Methods). When applying the rarefaction analysis on a single duplicate, we found that the rarefaction curve did not tend to an asymptote (*Figure 2A),* meaning that a continuous increase of sequence depth increases V_H_ sequence diversity. This is probably a result of the errors that are continually introduced during BCR-Seq, which precludes our ability to reach saturation of the rarefaction curve. In contrast, with the technical duplicate approach, the rarefaction curve of the V_H_ sequences (that are shared between the two duplicates) did saturate, thereby indicating that the sequencing depth did indeed suffice and that errors were sufficiently removed from the datasets (*Figure 2B-C, Supplementary Figure S1*). Subsequent application of the Spearman rank-order correlation on the BCR-Seq duplicates showed that antibody V_H_ sequence frequencies were correlated between duplicates (*Figure 2D-E, Supplementary Figure S2*). Rarefaction analysis of the validated sequences that were shared between the duplicates, from all tissues, demonstrated that the curve reached an asymptote, with 60-80% of the reads accounting for 99% of the possible diversity in the sample (*Figure 2F*). We thus showed that the technical duplicate approach was efficient in reducing the rates of errors introduced during BCR-Seq and that sufficient sequencing coverage was reached with high reproducibility. Details regarding the number of V_H_ sequences obtained in each computational step are summarized in *Supplementary Table S2.*

**Figure 2.**
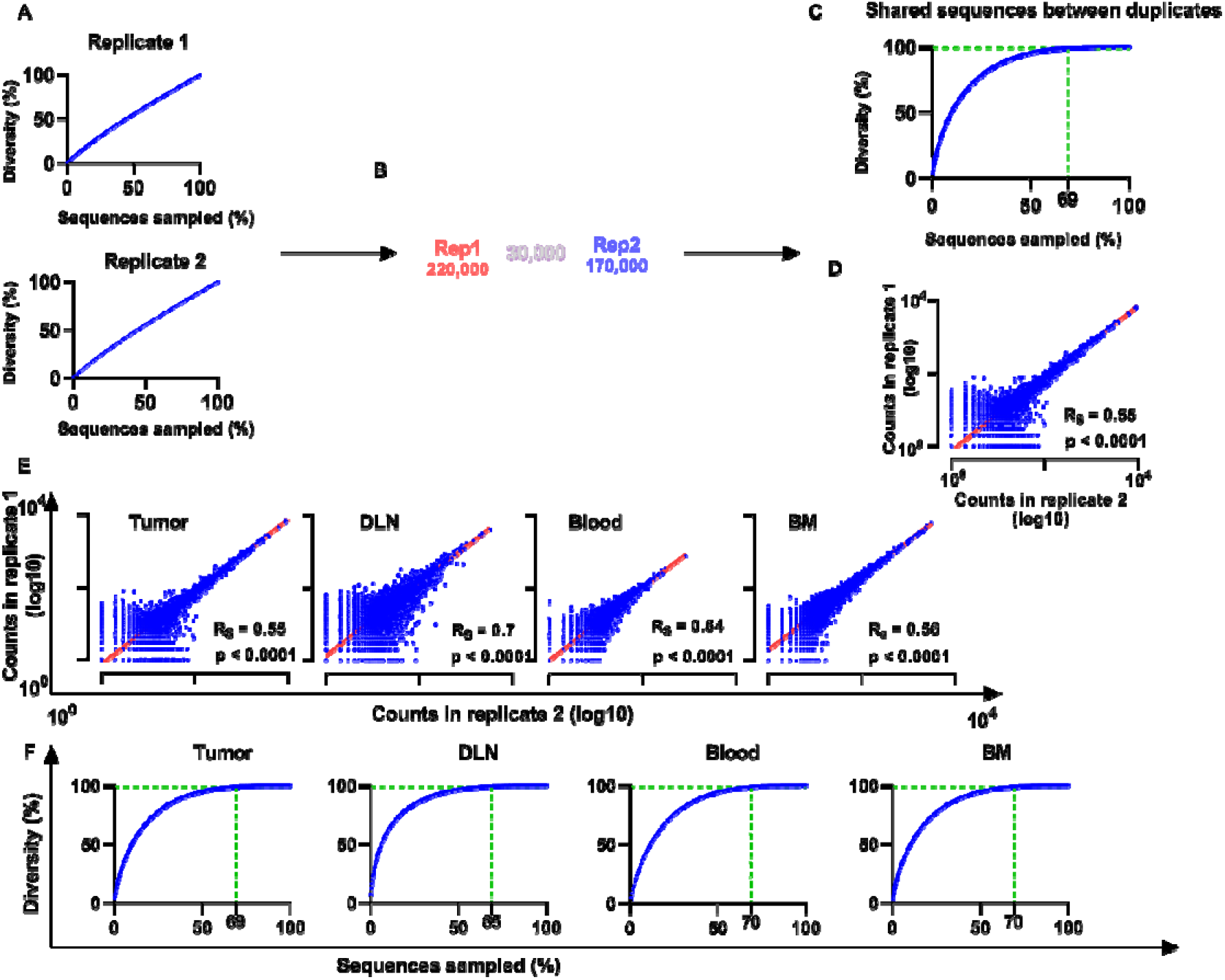
Duplicate approach for removal of errors from BCR-Seq data. **(A)** Rarefaction curves for each duplicate originating from the same sample. X-axis represents the percent of sequences out of the total number of sequences in the sample, and Y-axis represents diversity of V_H_ sequences and, hence, the percent of unique V_H_ sequences out of the total number of V_H_ sequences that were sampled. (**B**) Venn diagram of the duplicates, showing an example of the number of sequences in each duplicate and the number of shared sequences between duplicates. (**C**) Rarefaction curve for the shared sequences as identified in both technical duplicates. The vertical dashed line indicates the percent of sequences in which diversity reaches 99%. (**D**) Correlation between duplicates. Transcript counts for each V_H_ sequence were correlated between duplicates. X and Y axes are on a logarithmic scale and represent the transcript counts for each sequence. Each sequence is represented by a blue dot. (**E**) Correlation between library duplicates from all tissues. (**F**) Rarefaction curves for sequences from all tissues. Rs – Spearman rank-order coefficient; p-values are indicated. BM – bone marrow; DLN – draining lymph node.

It should be noted that for all the repertoire analyses V_H_ sequences from the four treated mice were pooled, as were the V_H_ sequences from the two untreated mice, and that individual datasets for each mouse were kept separate when describing common clones between tissues.

### TIL-Bs exhibit high clonal polarization and low clonal diversity

To determine the clonality of B cells in all tissues, we clustered the B cells on the basis of their identical V and J gene segments and 100% identity of the CDRH3 regions (nucleotide identity). The B cell clonal architecture was described in terms of two parameters: clonal polarization and clonal diversity. For the first parameter, high clonal polarization was defined as a state in which a few B cell clones contribute to the vast majority of antibody reads in a particular sample (*Figure 3A*). Thus, to quantify the clonal polarization level in each tissue, we calculated the number of B cell clones that gave rise to 80% of all immunoglobulin reads in a specific tissue. We found that, on average, 20 intra-tumoral immunoglobulin clonotypes dominated the B cell response in the TME, while in the other tissues the number of clones that contributed to 80% of the reads was higher than that in the TME by 2-3 orders of magnitude (*Figure 3B*).

**Figure 3.**
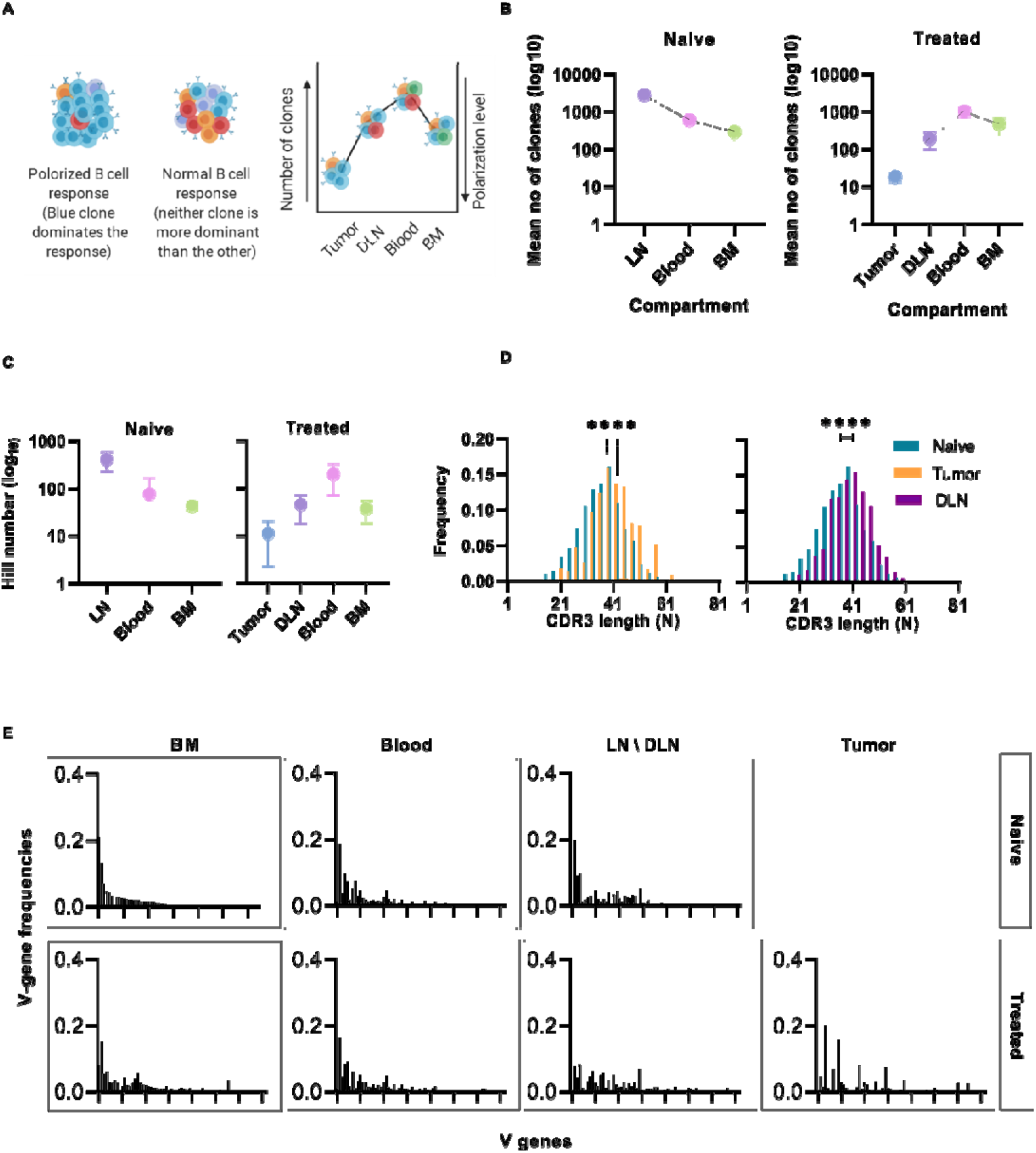
Clonal analysis, CDRH3 length distribution, and V gene usage of B cells isolated from different tissues. **(A)** Schematic representation of clonal polarization. Each color represents a single clone. As the number of clones increases, the polarization decreases. (**B**) Polarization curve; the mean number of clones (log10) that contribute to 80% of all B cell reads in each tissue is shown on the y-axis. The lower the value, the higher the polarization level of the B cell response. Error bars represent the standard error of the mean (SEM). (**C**) Hill number index (log10) – used as a measure of clonal diversity – was calculated for all clones in each tissue. Error bars represent standard deviation. **(D)** CDRH3 length distribution of intra-tumoral and DLN V_H_ reads. The median CDRH3 length of clones from the tumor or the DLNs of treated mice (orange/purple bars) was compared to that of naïve clones (red bars). Statistical significance was determined by using an nonparametric, unpaired, two-sided, Mann-Whitney t-test. For both the DLNs and the tumor, the p-value <0.05 was considered statistically significant. (**E**) Bar plots of V gene usage frequency in each tissue. All V genes were sorted according to the frequency of V genes vs. the naïve B cells (IgM^+^ B cells from the bone narrow of naïve mice). Naïve – naïve mice, Treated – tumor-bearing mice. two-sided, **** p ≤ 0.0001.

The second parameter that we calculated was clonal diversity, which reflects the number of unique B cell clones present in each tissue. This parameter can be used as an indicative measure to determine whether the nature of the response is oligoclonal or polyclonal. The Hill number diversity index^37^ indicated that the clones in the tumors were significantly less diverse than those in other tissues, a finding that is in keeping with the data for clonal polarization (*Figure 3C*).

The low diversity and the high polarization level in the TIL-B response suggests that B cell clonal proliferation occurs within the tumor tissue in our TNBC model. As expected, the B cell response in the DLNs, which are adjacent to the primary tumor, also showed a high clonal polarization level (compared to the “naïve” lymph nodes) (*Figure 3B*) and a low Hill number index (*Figure 3C*).

We then analyzed the length distribution of the CDRH3 to be used as an indicative measure for the positive clonal selection process that may take place in the tumor or the adjacent DLNs. The average CDRH3 length in the TIL-B clones was significantly longer than the CDRH3 length of B cell clones in the bone marrow of naïve mice. The same bias in the CDRH3 length was observed in DLN-derived B cells (*Figure 3D*). Bias in the CDRH3 length distribution may suggest that TIL-Bs are subjected to a positive selection pressure as part of the affinity maturation stage. Thus, subsets of B cells in the TME and DLNs may be considered as tumor-reactive B cell clonotypes that were possibly activated following engagement with the tumor cells or TAAs.

An antibody repertoire can be described in terms of the frequencies with which it uses the gene segments (V(D)J), particularly the V gene segment, as it is the longest and most diverse segment^38^. V gene usage in naïve B cell subsets is determined by a semi-random process, resulting in the association of the V-D-J segments during gene rearrangement as part of B cell development^39^. Thus, we posit that the V gene usage of IgM^+^ B cells isolated from the bone marrow of naïve mice can be used to determine the “basal” V gene usage, as previously shown^40^. The V gene usage in all tissues was sorted according to the descending frequencies vs the “basal” V gene usage. We found that the V gene usage signatures in the TIL-B and DLN clones were substantially different to the “basal” signature (*Figure 3E*). This perturbation of V gene usage can be considered as an indicator of the clonal selection process that occurs following an antigenic encounter.

### TIL-Bs are dominated by IgM that exhibits high SHM rates

Our data revealed that in both naïve and treated mice, IgM was the dominant isotype in the bone marrow B cells, while a small fraction of B cells were IgG^+^ (75% IgM and 20% IgG, *Figure 4A*). We observed a similar distribution when analyzing murine B cells derived from the blood; in all the mice, treated and non-treated, the dominant isotype was IgM (96%), reflecting antibodies encoded by either naïve B cells or non-CSR memory B cells (*Figure 4A*). Upon reaching the lymph nodes, B cells encounter antigens and undergo affinity maturation and CSR in germinal centers^45–47^. As expected, the IgG/IgM ratio encoded by B cells in the DLNs of the treated mice was higher than that in the lymph nodes of the naïve mice. Finally, TIL-Bs were mostly dominated by IgM and to a lesser extent by IgG (*Figure 4A*).

**Figure 4.**
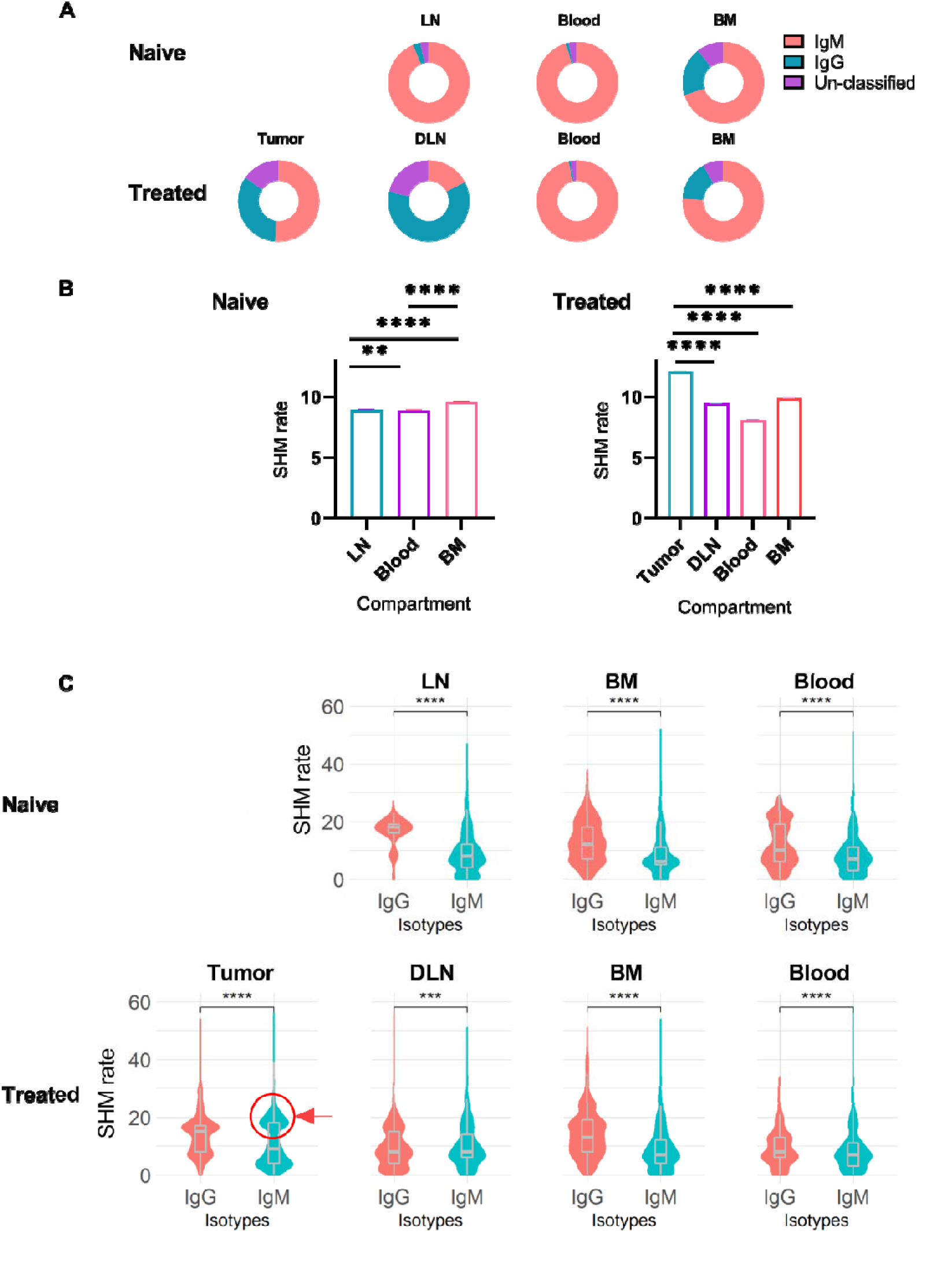
Isotype distribution and SHM rate in B cells isolated from different tissues. **(A)** Doughnut chart for the distribution of immunoglobulin isotypes encoded by B cells isolated from different tissues in treated and untreated mice. (**B**) Bar plots of SHM rate identified in the V_H_ regions of B cells isolated from each tissue. Error bars represent the standard error of the mean (SEM). (**C**) Violin charts showing the distributions of the SHM rate within each immunoglobulin isotype across tissues. The violin curves show the density of values; median, lower and upper quartile values are indicated. The red circle and arrow highlight the IgM^+^ B cell population that exhibits high SHM rates. Statistical significance for **B** and **C** was evaluated with the nonparametric, unpaired Mann-Whitney t-test (two-sided; **** p ≤ 0.0001, *** p ≤ 0.001, ** p ≤ 0.01). Naïve – naïve mice, Treated – tumor-bearing mice.

To further explore repertoire measures that are indicative of the activation state of B cells, we analyzed the SHM rates within each tissue for the different isotypes. We found that the SHM rate in the TIL-B compartment (mean of 12 mutations) was significantly higher than that in the DLNs, bone marrow and blood. Surprisingly, although the SHM rate was high, TIL-Bs were dominated by IgM (*Figure 4A-B*). To further investigate this observation, we compared the SHM rates between isotypes in all tissues. As expected, IgG^+^ B cells exhibited a significantly higher SHM rate than IgM^+^ B cells, since IgG^+^ B cells are considered to be activated B cells and have thus already gone through the process of affinity maturation and CSR. In addition, examining the SHM rate distribution in the IgG^+^ B cell subset revealed several levels of SHM, representing B cells that are at different affinity maturation stages (*Figure 4C*). Further examination of the SHM rate distribution in IgM^+^ B cells revealed a unimodal distribution (single dominant peak) in all tissues, except in the tumor, where the distribution was bimodal (one peak with a low SHM rate and the other with a high SHM rate, *Figure 4C*). The second peak in the IgM^+^ TIL-B subset (highlighted with a circle in *Figure 4C*) represents B cells that have undergone affinity maturation, but without CSR. A possible explanation for this interesting observation is that we harvested the TIL-Bs before they had had sufficient time to complete the CSR process. Another possible explanation is that CSR signaling was impaired due to tumorigenic activity in the TME.

### TIL-B clonal distribution across tissues

We identified, on average, 1250 B cell clones in the tumor, while the other tissues exhibited higher numbers of B cell clones (DLNs = 4700, blood = 5550, bone marrow = 12,110, *Supplementary Table S3*). To track the TIL-B clonal distribution in DLNs, blood and bone marrow, we identified clones that are common to the tumor and to any one of the three other tissue types. We thus defined clones that appear in the tumor and in at least one additional tissue as common clones and examined their frequency in each tissue (*Figure 5A;* details regarding the number of common clones are listed in *Supplementary Table S4*). The analysis of common clones was carried out for each mouse separately, because it is rare to find the common clones across different individuals. Thus, to obtain insights as to whether the B cell repertoire has convergence attributes, namely, similar repertoire measures of common clones between different mice, we looked at each mouse as a separate individual.

**Figure 5.**
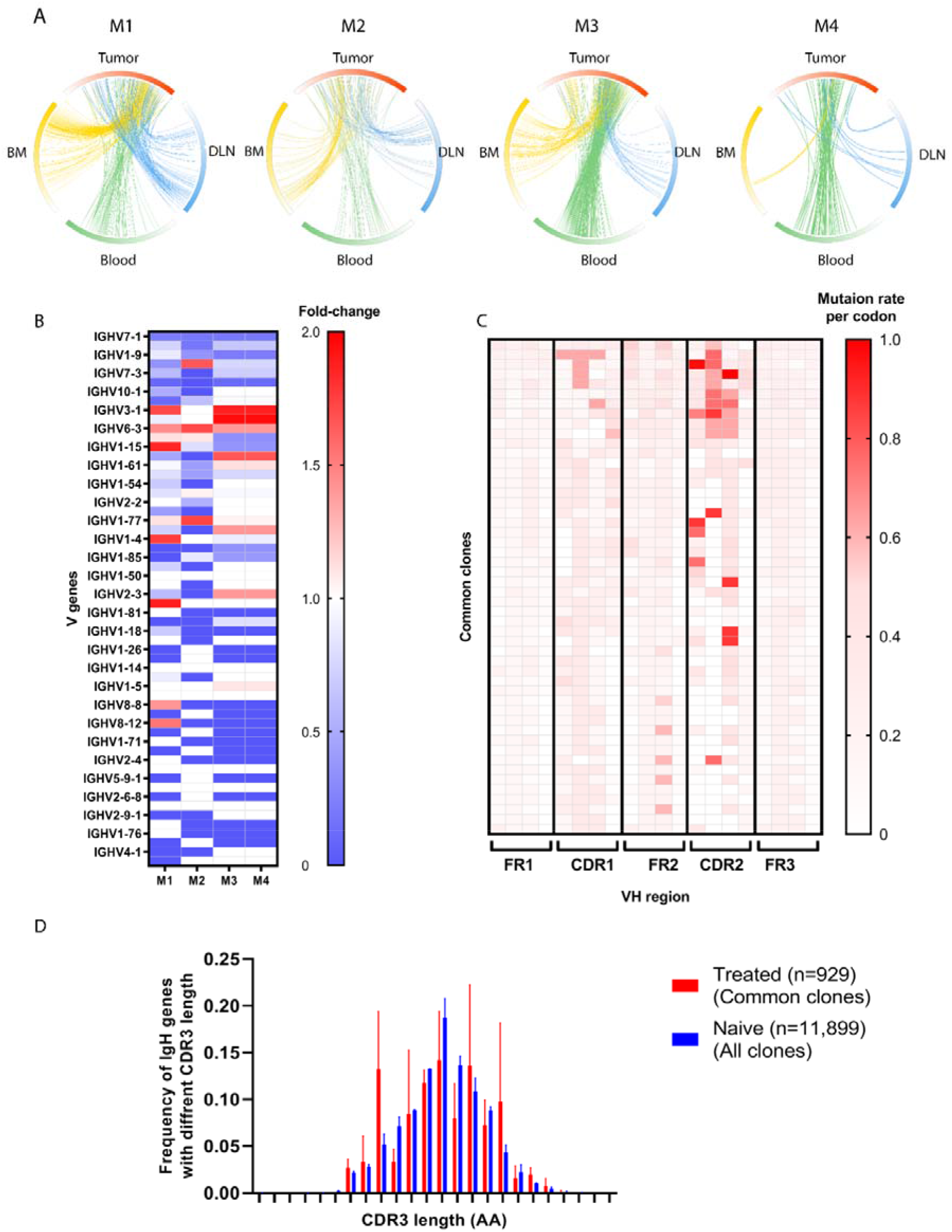
Antibody repertoire measures for the common clones. **(A)** Circos plots showing the clonal distribution of TIL-Bs between tissues. M1-4 indicate a circos plot for each mouse. The circos plot arcs are colored by gradients that are in accordance with the frequency of the clone within each tissue. Each connecting line represents a clone that is common between two tissues. Red – tumor, blue – DLN, green – blood, yellow – bone marrow. (**B**) Heat map showing fold-change of V gene usage frequency between naïve and treated mice. Each row represents a mouse (M1-4) and each column, a V gene. Red gradient represents an increase in the frequency of the V gene in the treated mice, while the blue gradient represents an increase in the frequency of the V gene in the naïve mice. (**C**) Heat map of the SHM rate per codon for each V_H_ region in the common clones. The red color indicates a higher SHM rate; each row represents one common clone; and each column represents a mouse. The heat map includes only the first 50 dominant common clones from each mouse. (**D**) The distribution of V_H_ genes with different CDRH3 lengths is shown in the bar graph. The average CDRH3 length of all the common clones for the treated mice (red bars) was compared to that for the naïve clones (blue bars). Statistical significance was determined by using an unpaired t-test. The p-value is p = 0.4939, which is considered to be not statistically significant (not shown). Error bars represent the standard deviation.

Among the B cell clones observed in the tumor, 20% were designated as common clones shared between the tumor and other tissues, suggesting that these clones had migrated to/from the TME. Although those common clones comprised only 20% of total clones in the TME, their V_H_ reads accounted for approximately 80% of all V_H_ reads in the TME.

We then set out to identify whether there were over-represented V genes in those common clones, namely, the clones that had been subjected to positive selection during affinity maturation, which skewed the V gene usage signature. A comparison of the semirandom V gene usage, as observed in the naïve tissue, with the V gene usage observed in the common clonotypes (*Figure 5B*) revealed 14 V genes that were over-represented in the treated mice, with six of them being common to at least two mice. The *IGHV6-3* V gene was over-represented in all four mice. These over-represented V genes may suggest that a converging process may have occurred in the common clones as a result of engagement with the tumor or TAAs in the TME.

In addition, we examined whether SHM patterns were introduced preferentially along the V_H_ regions (FRs and CDRs) within the common clonotypes. As expected, CDRH1-2 exhibited increased rates of SHM compared to the adjacent FRs (*Figure 5C*).

Next, we examined whether the distribution of CDRH3 length was skewed in the common clones versus all clones identified in the naïve mice (*Figure 5D*). While the median CDRH3 length of both the common clones and the naïve clones was 13 AA (n = 930, n = 11995, respectively), the CDRH3 length distribution of the common clonotype did indeed show a distribution pattern that was different to that of the naïve B cells.

Lastly, mice produce four IgG subclasses, namely, IgG1, IgG2A, IgG2B and IgG3. Specifically, it has been suggested that a low IgG1/IgG2A frequency ratio reflects the dominance of Th1 responses over Th2 responses. The production of IgG1 rises in response to the Th2 cytokine IL-4, and the production of IgG2A rises in response to the Th1 cytokine IFN^^41,42^ To examine any alterations in the ratio of IgG1 to IgG2A frequencies, as found in the common clones, we calculated the IgG1/IgG2A ratio in B cells from the bone marrow, blood and lymph nodes of naïve mice and compared it to the ratio as found in the tumor common clones in each of the above tissue types in the tumor-bearing mice (i.e., the clones common to the tumor and the bone marrow, blood or DLN). We found that IgG1/Ig2A ratio in the common clones was, on average, twofold lower than the ratio in the naïve mice (*Supplementary Figure S3*).

## Discussion

Reports on the role of TIL-Bs in breast cancer are contradictory, with some studies suggesting that the infiltration of B cells into the TME is associated with a poor prognosis^43,44^, but others suggesting that they correlate with improved prognosis^45^. Regardless of the specific role of B cells in the TME, it remains an open question whether B cells in the TME are reactive towards tumor cells and whether they traffic to various body compartments. To address this question, we used antibody repertoire measures to profile and track TIL-Bs in a TNBC mouse model.

We utilized an experimental and computational platform to elucidate repertoire measures in four tissues of TNBC mice. First, we performed several quality assurance steps to make sure that the antibody sequences were of high quality and that the sequences obtained were reproducible. This was achieved by using experimental duplicates for all NGS libraries. This approach, accompanied by applying additional filters, resulted in high-quality antibody V_H_ sequence datasets. The resultant high-quality datasets were then further analyzed to identify repertoire measures with distinct signatures suggesting that TIL-Bs did indeed engage with tumor cells and/or TAAs in the TME.

This study thus provides evidence that B cell populations in the tumors and the DLNs of TNBC mice were enriched with clones that exhibit the characteristics of activated, affinity-matured B cells, as indicated by increased SHM rates and skewed V gene usage and CDRH3 length. Moreover, by comparing the clonality of TIL-Bs with that of B cells in the DLNs, bone marrow and blood, we found that TIL-Bs exhibited low clonal diversity and high clonal polarization, namely, a limited number of distinct B cell clones were prevalent and these accounted for the greater part of the B cell response in the TME. Of note, this observation of polarized clonality in the TME has been described in other breast cancer studies^46,47^ as well as in studies of ovarian cancer^48^ and melanoma^26,49^, which also reported that a small number of clones dominate the B cell response in the TME. These studies taken together suggest that the B cells in the TME are constantly challenged by TAAs or by the tumor cells themselves. Moreover, the clonality of the B cells in the adjacent DLNs was also polarized, although to a lesser extent. With regard to isotype distribution, IgM^+^ B cells in the bone marrow are immature and/or naïve-mature B cells that have developed from hematopoietic stem cells, and therefore predominantly express the IgM isotype on their surface, whereas bone marrow IgG^+^ B cells can be considered as antigen-specific, terminally differentiated B cells, which home to the bone marrow following affinity maturation^50,51^. IgM^+^ naïve B cells migrate from the bone marrow to secondary lymphoid organs via the blood circulation^50^, and it has been reported that, in humans, circulating B cells in the blood comprise mostly of naïve B cells (IgM^+^, 70%) and memory B cells that are either class-switched or non-class switched (30%)^52^. As expected, in our study, the dominant isotype in the blood was found to be IgM, but the dominant isotype in the DLNs was skewed towards IgG. Surprisingly, the dominant isotype in the TME was found to be IgM, rather than IgG, and more importantly, we identified a population of non-class switched IgM^+^ B cells in the TME that exhibited high SHM rates; this population may represent B cells that have undergone affinity maturation even though their ability to carry out CSR has been inhibited. It is generally accepted that CSR and SHM take place in germinal centers and that these events are independent, with neither one being a prerequisite for the other^53^. However, recent work re-visiting this question showed that CSR is triggered before differentiation into germinal centers and that the vast majority of CSR events occur before the onset of SHM^54,55^. Nonetheless, whether CSR starts early in the process of B cell activation or not, the finding of a subset of highly mutated IgM^+^ B cells is intriguing in that it gives rise to the possibility that the TME generates inhibitory signals that prevent the full development of TIL-Bs and that hamper their ability to differentiate and encode for antibodies with appropriate effector functions.

We also analyzed the trafficking of TIL-Bs to other body tissues (DLNs, blood and bone marrow) by identifying common clonotypes. We found that an average of 20% of TIL-B clones are common to the tumor and at least one other tissue and that these common clones are highly prevalent in the TME. Moreover, V gene usage of the common clones was skewed towards several dominant V genes (some of them found in two of the four mice), suggesting that these clones had been subject to positive selection. Furthermore, as expected, the SHM in the common B cell clones occurred preferentially in the CDRs, which are unstructured loops in antibody binding sites^56^. Interestingly, the decrease in the IgG1/IgG2A ratio in the IgG subset of the common clones suggests that TIL-B cells encode mostly to IgG2A antibodies that fix complement^57^ and bind to all activating FcγRs, rather than to the IgG1 subclass that does not fix complement and that binds well only to inhibitory FcγRIIbs^58^.

The identification of TIL-Bs exhibiting reactive repertoire measures (tumor-reactive B cells or TRBCs) holds great potential for application in future studies. For example, the combination of antibody repertoire analysis and global gene expression profiling will allow further explorations of the possible cross-talk between TRBCs and the TME, thus facilitating an understanding of the role of B cells in the TME, with direct implications for the development of advanced immunotherapies.

## Data availability

Raw data can be accessed from the NCBI sequence read archive (BioProject ID: PRJNA699402).

## Code availability

The developed R codes for antibody repertoire analysis can be access upon request from: https://github.com/ligalaizik/Antibody-repertoire-analysis-R-code-

## Acknowledgements

The study was partially supported by

## Methods

### In vivo tumor models and control mice

TNBC 4T1 cells were purchased from the ATCC. BALB/C mice were injected subcutaneously into mammary fat-pad number five with 2×10^5^ 4T1 cells suspended in 30 μl of DMEM (Gibco, Thermo Fisher Scientific). The induced tumors were monitored by measuring their size twice a week using calipers. After 23 days, mice were sacrificed, and B cells were isolated from four tissue types, namely, bone marrow, blood, DLNs, and tumors. For the control mice, three tissue types were collected – bone marrow, blood, and lymph nodes.

### B cell isolation

All tissue preparations were taken at the same time from each mouse (after euthanasia by CO_2_ inhalation). For isolation of B cells from the lymph nodes, the lymph nodes were removed from the euthanized mice and mashed through a 70-μM cell strainer. Cells were then washed by centrifugation at 600 rcf for 5 min at 4–8 □. For isolation of TIL-Bs, tumors were enzymatically digested with 2,000 U/mL of DNase I and 2 mg/mL of collagenase IV (both from Sigma Aldrich, Merck, Israel) in HBSS for 30 min 37 □. with a magnetic stirrer (200 rpm). Cells were then washed by centrifugation at 600 rcf for 5 min at 4–8 □.. For obtaining B cells from peripheral blood, peripheral blood was collected by perfusion into sodium heparin-coated vacuum tubes before 1:1 dilution in HBSS supplemented with 2% FBS and 5 mM EDTA. For isolation of B cells from bone marrow, the femurs and tibias were removed from the tumor-bearing mice, ground using pestle and mortar in RPMI 1640 (Gibco, Thermo Fisher Scientific), filtered through a 70-μM cell strainer, and washed twice with RPMI 1640. Lymphocytes from all tissues were enriched on a Histopaque-1077 Hybrid-Max (Sigma Aldrich, Merck, Israel) density gradient medium, and the collected cells were washed twice with a complete RPMI 1640. For all tissues, cells were then incubated with a mixture of anti-IgG, anti-IgM, anti-CD138 magnetic beads (MojoSort™ Nanobeads, BioLegend, Carlsbad, CA) according to the manufacturer’s instructions.

### Amplification of V_H_ repertoires from B cells (BCR-Seq)

Total RNA was extracted from enriched B cells using the RNeasy Micro Kit (Qiagen, cat no. 74004) according to the manufacturer’s protocol. RNA concentration was determined on Epoch™ Microplate Spectrophotometer (BioTek). RNA was stored at −80 °C. Later, first-strand cDNA was synthesized using SuperScript™ III First-Strand Synthesis System (Thermo Scientific Cat no. 18080051) with 200 ng RNA as the template and Oligo (dT) primers (Thermo Scientific), according to the manufacturer’s protocol. After cDNA synthesis, PCR amplification of the variable heavy Ig genes was performed using a set of 19 forward primers with the gene-specific regions annealing to framework 1 of the VDJ-region and two reverse primers with the gene-specific region binding to the IgG and IgM constant regions (see primer list in *supplementary, Table S3*). Primers had overhang nucleotides to facilitate Illumina adaptor addition during the second PCR. PCR reactions were carried out using FastStart™ High Fidelity DNA polymerase (Sigma-Aldrich Cat no. 3553400001) in a reaction volume of 250 μl using 10 μl of the cDNA product as a template, and setting the following conditions as standard: 95 °C for 3 min; 4 cycles of 95 °C for 30 s, 50 °C for 30 s, 68 °C for 1 min; 4 cycles of 95 °C for 30 s, 55 °C for 30 s, 68 °C for 1 min; 20 cycles of 95 °C for 30 s, 63 °C for 30 s, 68 °C for 1 min; 68 °C for 7 min; 4 °C storage. PCR products were purified using Agencourt AMPure XP beads (Beckman Coulter, Cat no. A63881), according to the manufacturer’s protocol (ratio × 1.8 in favor of the beads). Recovered DNA products from the first PCR was applied to a second PCR amplification to attach Illumina adaptors to the amplified V_H_ genes using the primer extension method, as described previously^60^. Second PCR reactions were carried out using Phusion^®^ High-Fidelity DNA Polymerase (NEB) with the following cycling conditions: 95 °C denaturations for 3 min; 98 °C for 10 s, 40 °C for 30 s, and 72 °C for 30 s for two cycles; 98 °C for 10 s, 62 °C for 30 s, and 72 °C for 30 s for 7 cycles; and a final extension at 72 °C for 5 min. PCR products were subjected to 1% agarose DNA gel electrophoresis and gel-purified with Zymoclean™ Gel DNA Recovery Kit (Zymo Research) according to the manufacturer’s instructions. V_H_ library concentrations were measured using Qubit Flex Fluorometer (Thermo Fisher Scientific), and library quality was assessed using the 4200 TapeStation system (Agilent). DNA length of all libraries was between 540-570 bp.

### Replicate sequencing

Technical replicates (two per sample) of BCR-Seq libraries were prepared based on cDNA from each mouse/tissue. cDNA was split, and library preparation was performed in parallel with different Illumina indices as described above.

### Illumina sequencing, data analysis, and statistics

V_H_ libraries from sorted B cells were subjected to NGS on the MiSeq platform with the reagent kit V3 2 × 300 bp paired-end (Illumina), using an input concentration of 16 pM with 5% PhiX. Forward and reverse raw fastq files were paired and CDR3 and full-length VDJ regions of successfully paired sequences were annotated using MiXCR^35^ with default parameters. For downstream analyses, sequences were pre-processed and read-only retained based on the following criteria: (i) reads were shared between 2 replicates of the same library; (ii) both CDR3 and VDJ regions could be detected by IMGT; (iii) VDJ regions were present with a minimum abundance of 2 reads.

### Computational analysis

#### Rarefaction analysis

For the rarefaction analysis, V_H_ reads were randomly and repetitively sampled, and the number of species (unique V_H_ reads) was evaluated as a function of sample size^36,59^.

#### Duplicates correlation

For the correlation between the replicates of the same library (duplicates), the count of shared V_H_ reads between the two replicates was compared using Spearman’s rank correlation. Statistical significance was evaluated using Prism software.

#### Repertoire measures analysxis

All repertoire measures described in this study are a result of pooling of measures from all four treated mice, and for the naïve cohort, pooling of two naïve mice, with the exception of the circos plots and the V gene usage in *Figure 6.*

For the clonal analysis, we defined clonally related B cells as those sharing the same V_H_ and J_H_ gene groups and the same CDR3 nucleotide sequence. To assess clonal diversity, we applied the Hill number diversity index, based on Rényi’s definition^37^.

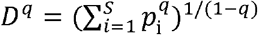

where *p_i_* is the frequency of each clone and *S* is the total number of clones. The *q* values represent weights; thus as *q* increases, clones with higher frequency have a stronger impact on the obtained Hill number index (in this case, *q*=2).

For the CDRH3 length analysis, we measured the number of nucleotides/amino acid starting at the end of FR3 (CAR amino acids) and ending with Y amino acid at the beginning of the J region.

For the V gene usage analysis, the number of unique V_H_ reads annotated to each V gene was quantified in each tissue. V gene usage of IgM-expressing B cell reads from the bone marrow of the two naïve mice was averaged, and then sorted from the most abundant V gene to the rarest V gene. Next, all V gene frequencies were sorted according to the naïve-bone marrow V gene order.

Isotype distribution plots are based on the number of unique V_H_ reads annotated to each isotype.

To examine the mutation level in each B cell repertoire, primer trimming was performed on all V_H_ reads before including nucleotide deletions, insertions, and substitutions, using in-house computational R scripts. All statistical tests in the manuscript were carried using R / GraphPad prism V9.0.2.

