## Supplemental for "Antibody repertoire analysis of tumor-infiltrating B cells reveals distinct signatures and distributions across tissues"

**Supplementary**

| PCR1-FW | GLUE + 5' Gene specific region |
| --- | --- |
| m-VH-glue-Fw1 | CCC TCC TTT AAT TCC CGA KGT RMA GCT TCA GGA GTC |
| m-VH-glue-Fw2 | CCC TCC TTT AAT TCC CGA GGT BCA GCT BCA GCA GTC |
| m-VH-glue-Fw3 | CCC TCC TTT AAT TCC CCA GGT GCA GCT GAA GSA STC |
| m-VH-glue-Fw4 | CCC TCC TTT AAT TCC CGA GGT CCA RCT GCA ACA RTC |
| m-VH-glue-Fw5 | CCC TCC TTT AAT TCC CCA GGT YCA GCT BCA GCA RTC |
| m-VH-glue-Fw6 | CCC TCC TTT AAT TCC CCA GGT YCA RCT GCA GCA GTC |
| m-VH-glue-Fw7 | CCC TCC TTT AAT TCC CCA GGT CCA CGT GAA GCA GTC |
| m-VH-glue-Fw8 | CCC TCC TTT AAT TCC CGA GGT GAA SST GGT GGA ATC |
| m-VH-glue-Fw9 | CCC TCC TTT AAT TCC CGA VGT GAW GYT GGT GGA GTC |
| m-VH-glue-Fw10 | CCC TCC TTT AAT TCC CGA GGT GCA GSK GGT GGA GTC |
| m-VH-glue-Fw11 | CCC TCC TTT AAT TCC CGA KGT GCA MCT GGT GGA GTC |
| m-VH-glue-Fw12 | CCC TCC TTT AAT TCC CGA GGT GAA GCT GAT GGA RTC |
| m-VH-glue-Fw13 | CCC TCC TTT AAT TCC CGA GGT GCA RCT TGT TGA GTC |
| m-VH-glue-Fw14 | CCC TCC TTT AAT TCC CGA RGT RAA GCT TCT CGA GTC |
| m-VH-glue-Fw15 | CCC TCC TTT AAT TCC CGA AGT GAA RST TGA GGA GTC |
| m-VH-glue-Fw16 | CCC TCC TTT AAT TCC CCA GGT TAC TCT RAA AGW GTS TG |
| m-VH-glue-Fw17 | CCC TCC TTT AAT TCC CCA GGT CCA ACT VCA GCA RCC |
| m-VH-glue-Fw18 | CCC TCC TTT AAT TCC CGA TGT GAA CTT GGA AGT GTC |
| m-VH-glue-Fw19 | CCC TCC TTT AAT TCC CGA GGT GAA GGT CAT CGA GTC |
| PCR1- REV | GLUE + 3' Gene specific region |
| m-IgMC-BC-glue-REV | GAG GAG AGA GAG AGA G CG AGG GGG AAG ACA TTT GGG |
| m-IgGall-BC-glue-REV | GAG GAG AGA GAG AGA G CC ARK GGA TAG ACH GAT GGG |
| PCR2-FW |  |
| PE-IgALL-Univ-FW | AAT GAT ACG GCG ACC ACC GAG ATC TAC ACT CTT TCC CTA CAC GAC GCT CTT CCG ATC TNN NNC CCT CCT TTA ATT CCC |
| PCR2-REV |  |
| YW23X_(reverse complement) PE-Idx-REV | CAA GCA GAA GAC GGC ATA CGA GAT **NNN NNN** GTG ACT GGA GTT CAG ACG TGT GCT CTT CCG ATC TNN NNG AGG AGA GAG AGA GAG |

**Table S1.** Primers used for primer extension BCR-Seq. PCR1 primers – FW: V_H_ gene-specific seqeunce (red) and GLUE for primer extension PCR (ligh blue); REV (reverse complement): GLUE (ligh blue), CH1 gene specific sequence (red). PCR2 primers – FW: GLUE for primer extentsion annealing (ligh blue), required diversity region (green), Universal trueSeq Illumina adaptor (purple); REV(reverse complement): GLUE (light blue), required diversity region (green) + T, Universal trueSeq Illumina adaptor.

| **Mouse** | **Tissue** | **Total raw reads** | **Productive reads** | **Unique reads** | **Shared in dupli­cated** | **Total reads**  **after filtration** |
| --- | --- | --- | --- | --- | --- | --- |
| **M1** | **Tumor** | 3,013,522 | 1,545,089 | 559,089 | 48,099 | 1,318,869 |
|  |  | 2,799,957 | 1,584,532 | 575,357 |  |  |
|  | **DLN** | 6,410,141 | 4,383,941 | 2,265,644 | 92,451 | 2,366,804 |
|  |  | 1,222,344 | 837,088 | 470,405 |  |  |
|  | **BM** | 5,311,991 | 3,838,182 | 1,803,273 | 184,023 | 3,001,259 |
|  |  | 2,764,938 | 1,989,530 | 1,051,214 |  |  |
|  | **PB** | 819,573 | 572,017 | 413,563 | 25,247 | 198,512 |
|  |  | 341,713 | 239,777 | 183,529 |  |  |
| **M2** | **Tumor** | 2,159,444 | 1236417 | 271922 | 21708 | 1,216,678 |
|  |  | 2,356,565 | 1495127 | 330478 |  |  |
|  | **PB** | 478,136 | 331,650 | 240,726 | 15,944 | 103,160 |
|  |  | 184,796 | 127,058 | 104,631 |  |  |
|  | **DLN** | 764,428 | 500940 | 342,847 | 28,840 | 233,689 |
|  |  | 887,901 | 590451 | 412,871 |  |  |
|  | **BM** | 1,362,027 | 952,835 | 526,383 | 81,425 | 875,626 |
|  |  | 1,559,131 | 1,104,857 | 587,970 |  |  |
| **M3** | **Tumor** | 1,044,197 | 648,208 | 220,884 | 28,072 | 680,448 |
|  |  | 758,064 | 469,970 | 167,227 |  |  |
|  | **PB** | 467,524 | 307,216 | 221,817 | 30,823 | 183,961 |
|  |  | 504,876 | 331,817 | 236,430 |  |  |
|  | **DLN** | 1,006,525 | 587,993 | 271,983 | 13,905 | 367,083 |
|  |  | 1,283,181 | 743,753 | 307,615 |  |  |
|  | **BM** | 725,273 | 484,223 | 277,649 | 51,213 | 653,939 |
|  |  | 1,418,660 | 965,328 | 500,111 |  |  |
| **M4** | **Tumor** | 172,148 | 121,997 | 30,847 | 7,995 | 228,652 |
|  |  | 275,603 | 190,847 | 43,671 |  |  |
|  | **PB** | 927,959 | 633,340 | 284,306 | 40,217 | 439,785 |
|  |  | 391,222 | 268,607 | 142,080 |  |  |
|  | **DLN** | 1,447,467 | 873,367 | 268,364 | 36849 | 926,849 |
|  |  | 1,248,512 | 749,077 | 249,827 |  |  |
|  | **BM** | 90,633 | 65,017 | 26,005 | 4168 | 76680 |
|  |  | 99,345 | 69,266 | 26,360 |  |  |
| **N1** | **LN** | 1,182,417 | 735,776 | 500,713 | 11722 | 45313 |
|  |  | 156147 | 91,531 | 82725 |  |  |
|  | **BM** | 1602662 | 1084211 | 755729 | 42223 | 356409 |
|  |  | 459945 | 307263 | 238084 |  |  |
|  | **Blood** | 1559966 | 1102181 | 612507 | 42566 | 510345 |
|  |  | 370609 | 242924 | 161998 |  |  |
| **N2** | **LN** | 640,750 | 414,668 | 337,538 | 22788 | 58056 |
|  |  | 564218 | 360,448 | 300636 |  |  |
|  | **BM** | 434917 | 300192 | 234853 | 17803 | 140258 |
|  |  | 485689 | 334271 | 263834 |  |  |
|  | **Blood** | 393680 | 263335 | 193244 | 23536 | 144656 |
|  |  | 383011 | 250325 | 187367 |  |  |

**Table S2. Summary of BCR-Seq data.** *Total raw reads*: reads obtained from the fastq files following paired-end alignment; *productive reads:* reads that were successfully aligned to the germline reference in IMGT; *exist in both duplicates:* unique V_H_ sequences that were shared between the technical duplicates; *total reads after filtration;* V_H_ sequences that were shared between technical duplicates (non-unique) and passed the filtration process.

T - treated, N - naïve.

| **Mouse #** | No. of clones | | | |
| --- | --- | --- | --- | --- |
|  | **Tumor** | **DLN** | **Blood** | **Bone marrow** |
| M1 | 1897 | 9212 | 4813 | 23,597 |
| M2 | 785 | 6418 | 3621 | 15790 |
| M3 | 1992 | 934 | 6766 | 8546 |
| M4 | 352 | 2289 | 7025 | 533 |
| Average | 1256.5 | 4713.25 | 5556.25 | 12116.5 |

**Table S3.** **Number of total B cell clones in the different tissues.**

| **Mouse #** | No. of TIL-B common clones | | |
| --- | --- | --- | --- |
|  | **DLN** | **Blood** | **Bone marrow** |
| M1 | 102 | 94 | 184 |
| M2 | 40 | 44 | 54 |
| M3 | 27 | 254 | 73 |
| M4 | 8 | 56 | 2 |

**Table S4. Number of common TIL-B clones.**

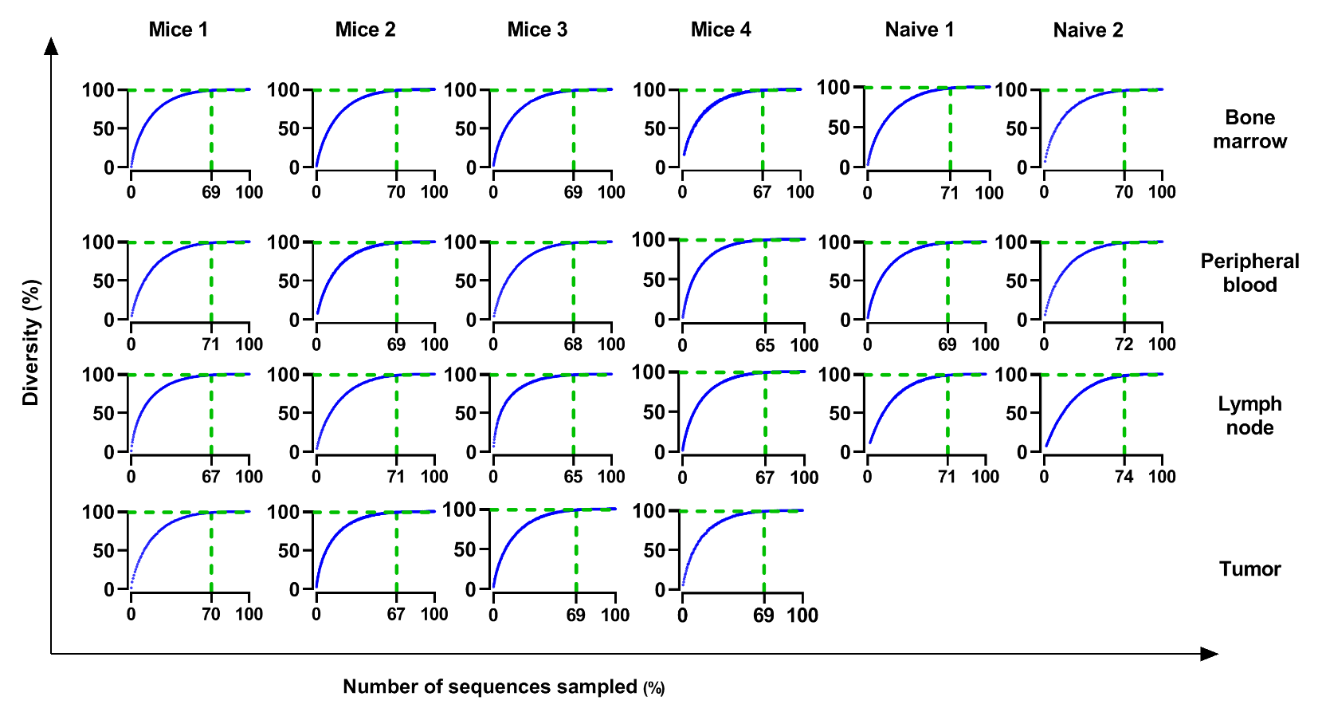

**Figure S1:** **Estimation of sequencing depth as a function of sampling.** Rarefaction curves represent sequence diversity (= number of unique sequences, Y-axis) as a function of the number of sequences sampled (X-axis). As the curve saturates, an increase in sample size does not affect sample diversity; therefore, the sequencing depth is adequate. The vertical dashed line indicates the number of sequences in which diversity reaches 99% of total diversity. X-axis units represent the fraction of the number of sequences in each sampling iteration out of the total number of sequences; Y-axis represents the fraction of unique seqeunces in each sampling iteration out of the total unique sequences.

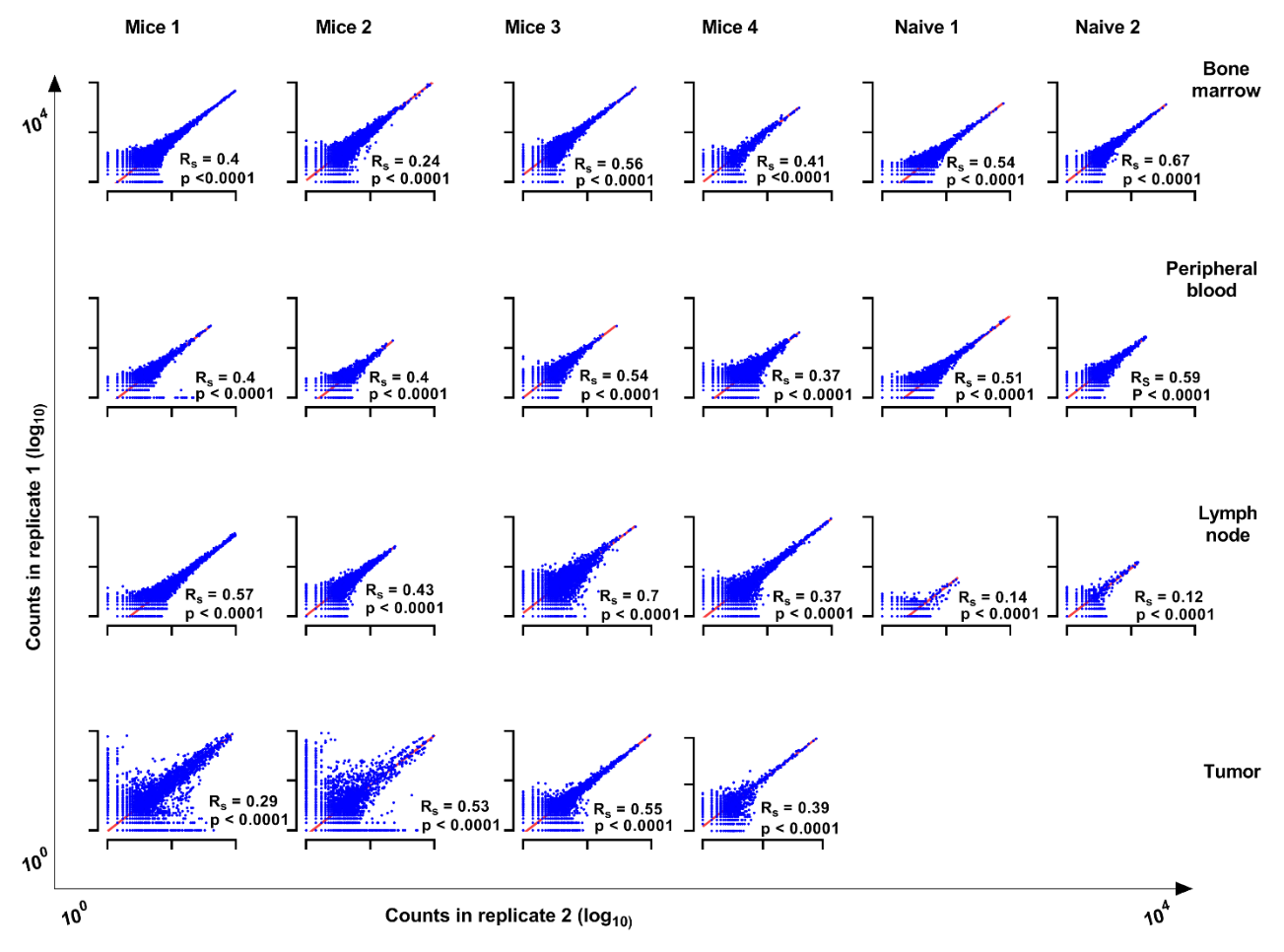

**Figure S2:** **Spearman's rank correlation of BCR-Seq duplicates**. Each dot represents a unique V_H_ sequence. The X and Y axes indicate the number of read counts for each V_H_ in the duplicates (logarithmic scale). R_S_ - Spearman's correlation coefficient, p – two tailed was considered significant if <0.05.

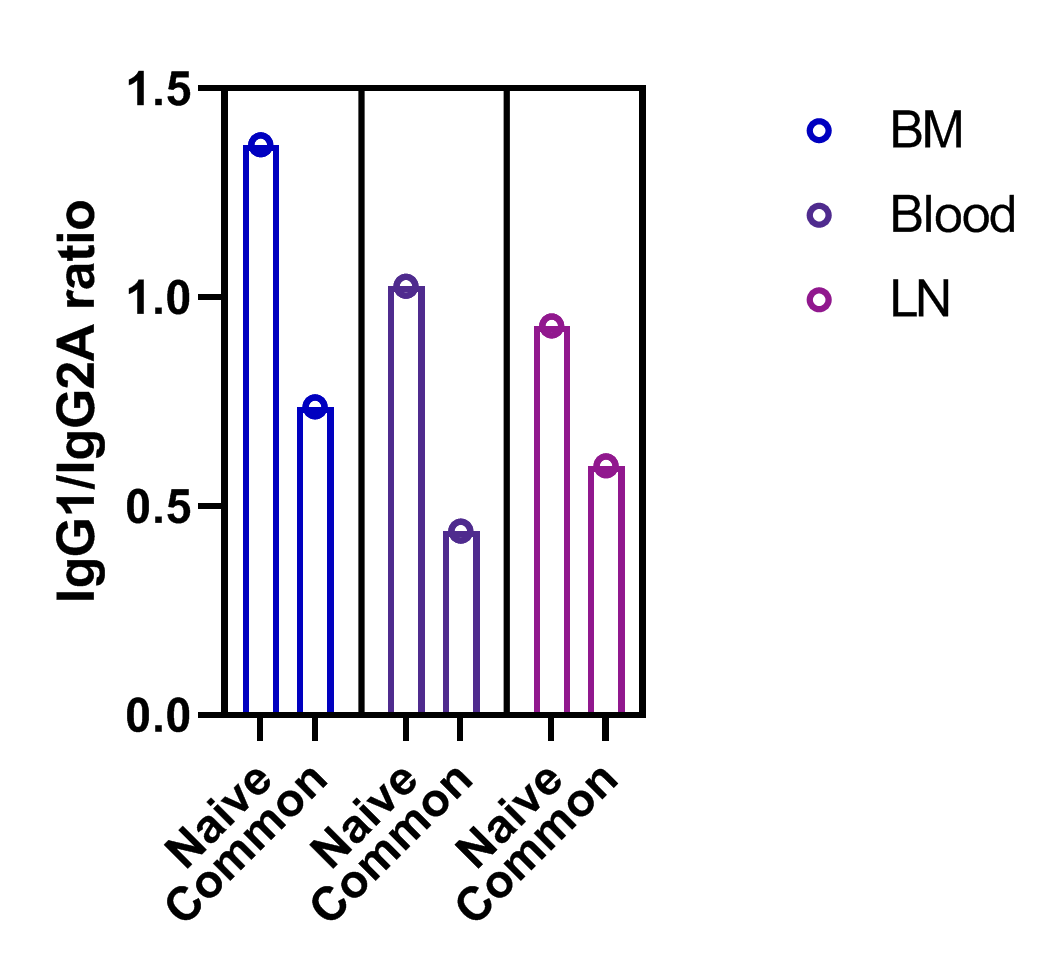

**Figure S3:** Calculated ratio between IgG1 and IgG2A subclasses. The ratio was calculated by dividing the relative frequency of antibodies with the IgG1 subclass by that with the IgG2A subclass as encoded by B cells in 3 tissue types from naïve mice and common clones to the tumor and the corresponding tissues in the treated mice (i.e., common to the tumor and bone marrow, blood or DLNs).
